# Allelic chromatin structure is a pervasive feature of imprinted domains and functions cooperatively with cis-acting long non-coding RNAs

**DOI:** 10.64898/2025.12.02.691962

**Authors:** Bongmin Bae, Katherine Gu, Daniel Loftus, Amanda Whipple

**Affiliations:** Department of Molecular and Cellular Biology, Harvard University, Cambridge, MA 02138, USA

## Abstract

Genomic imprinting is an epigenetic phenomenon in which genes exhibit restricted or biased expression from one allele according to parental origin. Imprinted gene expression plays crucial roles in embryonic growth and brain development. Higher-order chromatin structure has long been associated with gene regulation, particularly in the context of spatial enhancer-promoter interactions. Because imprinted genes exhibit allele-biased expression, it is compelling to ask whether differences in the three-dimensional organization of maternal and paternal genomes underlie this regulation. Using a capture Hi-C approach, we identified parental allele-specific higher-order chromatin structures across multiple imprinted domains in the mouse brain. These allele-specific structural features largely stem from annotated imprinting control regions (ICRs), concomitant with allele-specific binding of CTCF. The transcription start sites of active and inactive alleles of imprinted genes engage in distinct distal chromatin interactions that differ in number and in the epigenetic states of their contact regions. CRISPR interference (CRISPRi) screening identified a distal cis-regulatory element that modulates imprinted expression at the *Mest*-*Copg2* locus in neurons, with its regulatory activity closely linked to allele-specific chromatin interactions. Further investigation revealed that both a cis-acting long non-coding RNA and allele-specific enhancer-promoter architecture modulates *Mest*-*Copg2* imprinted expression in neurons. Together, this study highlights the interplay between chromatin structure and regulatory landscapes that modulate allele-specific expression of imprinted genes.

## Introduction

Three-dimensional (3D) genome organization in the nucleus spans multiple hierarchical scales and has been widely implicated in gene regulation (Dekker, 2008; Pombo & Dillon, 2015). At the compartment level, chromatin is partitioned into A and B compartments, which largely correspond to transcriptionally active and inactive regions, respectively (Lieberman-Aiden et al., 2009). At a finer scale, chromatin is organized into topologically associating domains (TADs) (Dixon et al., 2012; Nora et al., 2012), within which gene regulatory mechanisms, such as chromatin loop-mediated enhancer-promoter communication, take place (Dowen et al., 2014; Phillips-Cremins et al., 2013; Rao et al., 2014). TADs also serve to insulate regulatory effects between neighboring domains (Lupiáñez et al., 2015; Wendt et al., 2008), although recent evidence suggests that some regulatory crosstalk can occur across TAD boundaries (Balasubramanian et al., 2024). Allele-specific chromatin conformations have been examined in mouse and human cells, revealing several parent-of-origin-specific structures genome-wide, including at imprinted domains (Rao et al., 2014; Richer et al., 2023; Tan et al., 2021). Since genomic imprinting is an epigenetic phenomenon where imprinted gene expression is restricted to or biased toward a single parental allele, these observations provide a strong rationale for further dissecting the contribution of chromatin architecture to the regulation of imprinted expression.

Imprinting control regions (ICRs) are key regulatory elements that orchestrate imprinted gene expression across long genomic distances and are uniquely characterized by germline-inherited, parent-specific CpG methylation maintained throughout development (Geuns et al., 2003; Proudhon et al., 2012; Stöger et al., 1993; Tremblay et al., 1997). Functioning as gametic differentially methylated regions (gDMRs), ICRs direct the establishment of secondary, somatic DMRs (sDMRs) that regulate imprinted expression at a subset of gene promoters (Brandeis et al., 1993; Stöger et al., 1993). The DMRs can directly modulate transcriptional access to overlapping promoters, restricting expression to the unmethylated parental allele. However, some imprinted genes lack promoter-associated DMRs, indicating ICRs can also exert control through long-range regulatory mechanisms.

Two such mechanisms have been described: first, cis-repressive long non-coding RNAs (lncRNAs) and second, higher-order chromatin architecture. In many loci, unmethylated ICRs drive transcription of long noncoding RNAs (lncRNAs) that act in cis to repress neighboring imprinted genes. For instance, *Snhg14* lncRNA transcriptionally silences paternal *Ube3a* expression in neurons (Meng et al., 2013), while *Kcnq1ot1* lncRNA acts as a scaffold to recruit and spread repressive chromatin modifiers at neighboring genes (Mohammad et al., 2010; Pandey et al., 2008; Terranova et al., 2008). At other loci, ICRs regulate distally located imprinted genes by shaping higher-order chromatin architecture via methylation-sensitive binding of the CCCTC-binding factor (CTCF), a key organizer of chromatin loops (Prickett et al., 2013). This was first characterized at the *H19*-*Igf2* locus, where CTCF binding to the unmethylated ICR insulates the maternal *Igf2* promoter from its distal enhancer, permitting only paternal *Igf2* expression (Hark et al., 2000; Kurukuti et al., 2006; Szabó et al., 2004). Over the past decades, additional locus-specific studies have shown that parental allele-specific chromatin interactions between enhancers and promoters are a central mechanism of imprinted expression (Kurukuti et al., 2006; Llères et al., 2019; Loftus et al., 2023). Together, these findings provide a framework for understanding how distally located imprinted genes are regulated through allele-specific chromatin architecture.

However, a systematic examination of allele-specific chromatin architecture across imprinted domains is still lacking. Although recent technical advances have revealed sub-TAD-level allelic biases, few studies have functionally linked these structures to imprinted gene expression. Moreover, how the two prevailing regulatory paradigms – cis-acting lncRNAs and chromatin insulation – cooperate to establish and maintain imprinting remains poorly understood. Here, we generated high-resolution parental allele-specific chromatin contact maps across multiple imprinted domains in mouse brain. To identify functional cis-regulatory elements, we performed CRISPRi screens within these domains, uncovering a functional enhancer whose activity depends on allele-specific chromatin interactions. Focusing on the *Mest*-*Copg2* locus, we show that imprinted expression is dually regulated by a lncRNA and distal enhancer. A distal enhancer differentially activates paternal *Mest* and maternal *Copg2* expression through alternative structures, while the nuclear-enriched *Mest*-XL lncRNA selectively represses paternal *Copg2*. These findings underscore the critical role of 3D genome folding in coordinating enhancer-mediated and lncRNA-mediated mechanisms of allele-specific gene regulation.

## Results

### Imprinted domains exhibit pervasive parental allele-specific chromatin architectures

Given the characteristic mono-allelic expression pattern of imprinted genes, we first asked whether imprinted domains exhibit allele-specific chromatin structures. A recent study of spatial chromatin organization suggested the presence of allele-specific chromatin structures at a subset of imprinted domains in the mouse brain (Tan et al., 2021). To build upon these findings and conduct a higher-resolution interrogation of the relationship between allelic chromatin architecture and imprinted expression, we performed region capture Hi-C using cortical tissue from reciprocal F1 hybrid mice (BL6 x CAST and CAST x BL6). Genome-wide Hi-C libraries were enriched for ∼23 Mb region spanning eight imprinted domains, and each replicate was sequenced at a depth of 170-190 million reads (7-8 million reads per Mb). After applying HICUP quality control for mapping, ligation, and PCR duplication, the number of valid paired reads ranged between 10 to 17 million, corresponding to 5-9% of total sequencing reads (**Extended Figure 1A**).

Sequencing reads were then phased by parental origin using strain-specific single nucleotide polymorphisms (SNPs). Only reads overlapping informative SNPs can be assigned to a parental allele, so approximately 25% of valid reads were allelically informative. Among these, ∼23% represented cis-allelic interactions (either BL6-to-BL6 or CAST-to-CAST), while only 2% corresponded to trans-allelic interactions (BL6-to-CAST; **Extended Figure 1B**). This distribution is consistent with the expectation that each parental homolog predominantly occupies its own nuclear territory.

Using the cis-allelic read pairs, we generated parental allele-specific Hi-C contact matrices. Our capture Hi-C libraries provided sufficient complexity to assess chromatin proximity at 25, 10, and 5 kb resolutions (**Extended Figure 1C**). All eight imprinted domains captured in this study, namely *Peg10*-*Sgce*, *Mest*-*Copg2*, *Peg3*-*Usp29*, *H19*-*Igf2*, *Kcnq1-Kcnq1ot1*, *Snrpn*-*Peg12*, *Meg3*-*Dio3* and *Peg13*-*Kcnk9*, exhibited pronounced allele-specific chromatin structures (**Figure 1A-H**). The differential chromatin features observed in the capture Hi-C included discrete TADs (shown as triangular outline), architectural stripes reminiscence of slow loop extrusion (illustrated as diagonal lines), and prominent chromatin loops between distal loci (represented as strong dots along the diagonal axis and arc plot) (**Figure 1A-H**, left). Subtraction plots between the two parental alleles, as well as chromatin loops called by HICCUPS pipeline, demonstrated notable differences in chromatin architectures (**Figure 1A-H**, right). Collectively, these results revealed pervasive chromatin structural biases between parental alleles across imprinted domains in the mouse brain.

**Figure 1.**
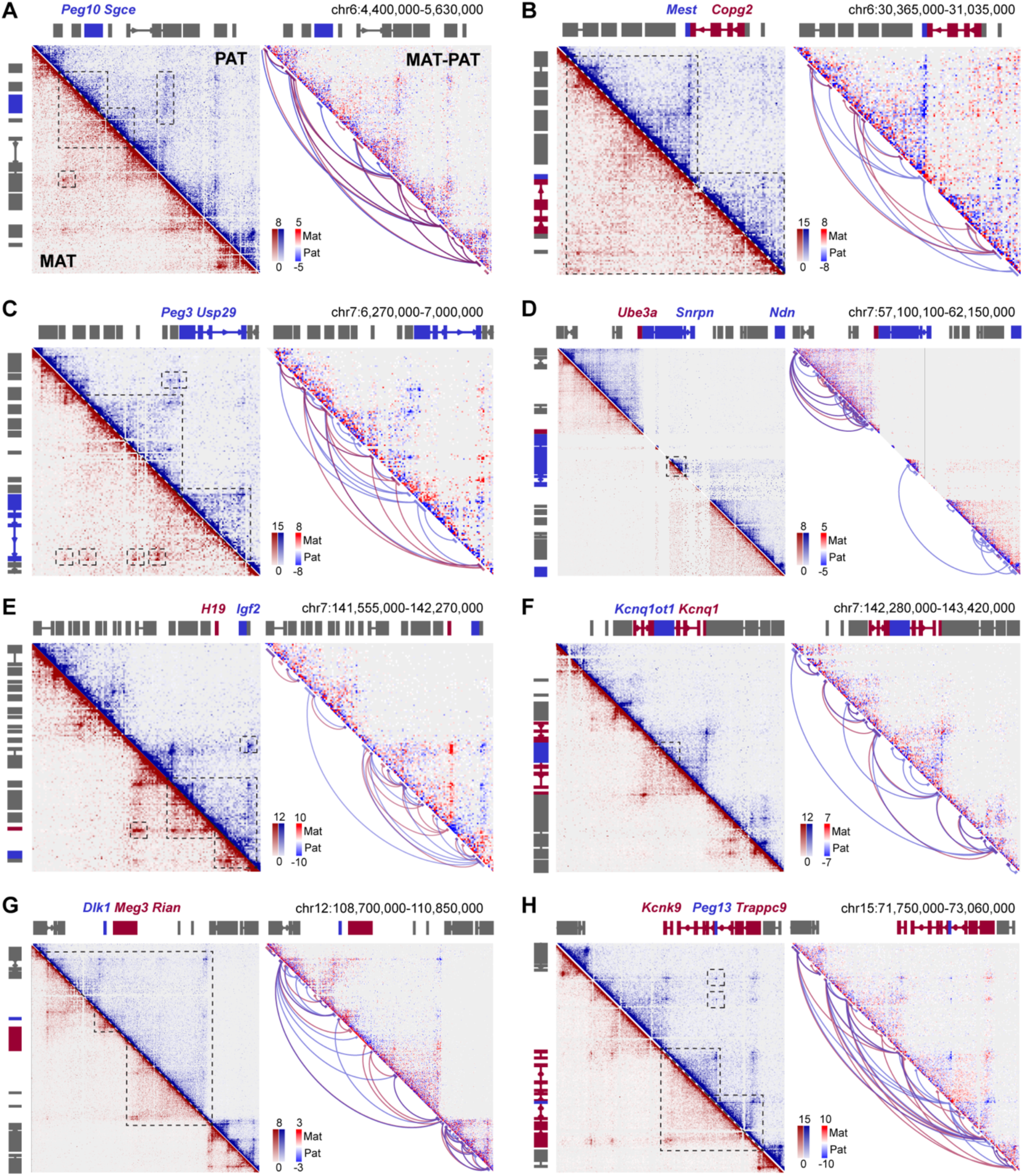
Imprinted domains exhibit pervasive parental allele-specific chromatin architectures. (**A**-**H**) *Left:* Capture Hi-C contact matrices constructed using allele-specific hybrid read pairs from mouse cortex at 5 kb resolution. Examples of chromatin structural features are highlighted in dashed outlines. Normalized using square root vanilla coverage (VC_SQRT) method; n = 4 merged replicates. *Right:* Allelic subtraction plots of contact matrices and arc plots showing significant loops called by HICCUPS pipeline. Maternal and paternal features are shown in red and blue, respectively.

### Methylation-sensitive CTCF binding at ICRs organizes allele-specific chromatin architecture

Given the central role of ICRs in regulating imprinted gene expression, we first assessed whether differential chromatin structures are anchored at DMRs. Allele-specific DMRs have been identified in the mouse brain, including both known gDMRs and sDMRs (**Extended Figure 2A**) (Xie et al., 2012). To determine whether these DMRs are associated with allele-specific chromatin organization, we performed pile-up contact analysis using our region capture Hi-C data. Chromatin interactions were assessed within 75 kb window upstream and downstream of each DMR.

Pile-up plots generated for methylated versus unmethylated alleles across 8 gDMRs, and 21 sDMRs revealed marked allele-specific differences in chromatin architecture. The methylated gDMR alleles displayed relatively uniform interaction patterns, whereas the unmethylated gDMR alleles exhibited local insulation of chromatin contacts across the DMRs, resulting in a checkerboard-like interaction pattern (**Figure 2A**). This allelic difference was less pronounced at sDMRs (**Figure 2B**). These results support the notion that unmethylated ICRs act as chromatin insulators or domain boundaries, restricting communication between adjacent genomic regions and thereby defining discrete regulatory domains.

**Figure 2.**
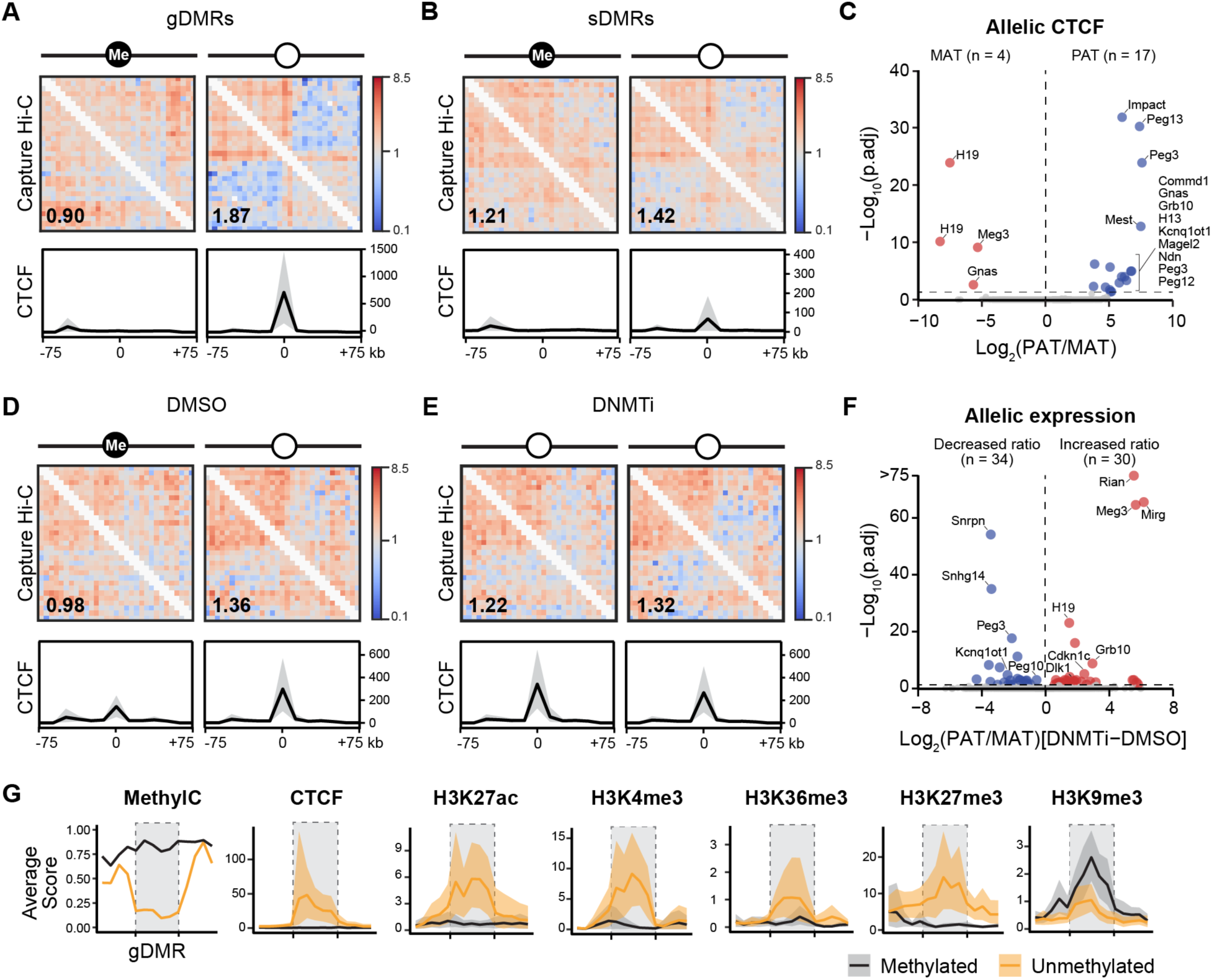
Methylation-sensitive CTCF binding at ICRs organizes allele-specific chromatin architecture. (**A-B**) *Top:* Pile-up density plots of cortex Capture Hi-C data generated by including 75 kb regions up- and downstream of each gDMR (**A**) and sDMR (**B**) separated by methylated vs unmethylated allele. Insulation score at DMR is shown in the bottom left corner of each plot. *Bottom:* Density plot of primary neuron CTCF tracks. (**C**) Volcano plot of paternally biased (blue) or maternally biased (red) CTCF binding peaks genome wide in mouse primary cortical neurons. (**D-E**) *Top:* Pile-up density plot of Capture Hi-C data from DMSO (**D**) or DNMTi-treated (**E**) mESCs across 75 kb regions up- and downstream of gDMRs separated by methylated vs unmethylated allele. Insulation score at DMR is shown in the bottom left corner of each plot. *Bottom:* Density plot of primary neuron CTCF tracks. n = 3 replicates per group. (**F**) Volcano plot of RNA-seq from DMSO vs DNMTi-treated mESCs illustrating changes in allelic expression ratio upon DNMTi treatment. p_adj_ is FDR corrected p-value of treatment x allele interaction. (**G**) Histone modification profiles from mouse brain on methylated vs unmethylated alleles across size-matched gDMR windows.

To assess whether CTCF binding at DMRs correlated with DNA methylation, we performed CTCF CUT&Tag in hybrid primary cortical neurons. We observed strong allele-specific CTCF binding at imprinted domains, with 17 paternally biased and 4 maternally biased CTCF peaks (**Figure 2C**). These peaks included our capture Hi-C target domains (*Peg10*-*Sgce*, *Mest*, *Peg3*-*Zim1*, *Peg12*, *H19*-*Igf2*, *Kcnq1ot1*, *Meg3*, and *Peg13)*, as well as additional imprinted loci (*H13, Gnas*, *Grb10*, *Commd1*, and *Impact)*, largely consistent with previous reports in mouse brain (Prickett et al., 2013). Although allelic CTCF binding at the *Snrpn* ICR did not meet the statistical threshold, potentially due to low SNP density in the region, it showed clear allelic bias (**Extended Figure 2B**). Allele-specific CTCF binding was preferentially enriched at unmethylated gDMRs and correlated with insulation effects (**Figure 2A,B**). However, the *Dlk1*-*Meg3* domain represents a notable variation: its unmethylated gDMR does not show strong allelic CTCF binding, and insulation instead arises from allele-specific CTCF occupancy at the sDMR at the *Meg3* promoter (**Extended Figure 2B**). This demonstrates that, in some imprinted regions, sDMRs can substitute for gDMRs as functional chromatin boundaries.

We then asked whether loss of DNA methylation could drive aberrant biallelic CTCF binding and disrupt allele-specific chromatin architecture at imprinted domains. We pharmacologically depleted DNA methylation in mouse embryonic stem cells (mESCs) using the DNA methyltransferase inhibitor GSK-3484862 (DNMTi), which induces DNMT1 degradation (Azevedo Portilho et al., 2021; Chen et al., 2023). Four days of treatment strongly reduced DNA methylation at ICRs (**Extended Figure 2C,D**) and led to loss of allelic chromatin structures. Specifically, the normally methylated allele showed increased insulation at the ICR following demethylation, accompanied by biallelic CTCF binding (**Figure 2D,E**).

Disruption of DNA methylation and chromatin architecture was associated with impaired imprinted expression (**Figure 2F**). Many imprinted genes became more biallelic: maternally expressed genes (e.g., *Meg3-Rian-Mirg*, *H19*, *Grb10*, and *Cdkn1c*) showed increased paternal/maternal expression ratios, consistent with reactivation of the normally silent paternal allele, whereas paternally expressed genes (e.g., *Snrpn*, *Sngh14*, *Peg3*, *Kcnq1ot1*, and *Peg10*) showed decreased paternal/maternal ratios, indicating reactivation of the maternal allele. The *Peg13*-*Kcnk9* domain was an exception, as imprinted expression at this locus remained unchanged, likely due to low expression of these genes in mESCs. Several other imprinted genes (*H13*, *Zdbf2*, *Ndn*, *Usp29*, *Mest*, and *Rtl1)* showed modest changes in allelic ratio that did not pass our statistical threshold.

The effects of DNMTi appear be cell-type specific: in primary neural precursor cells (NPCs), treatment produced only partial (∼50%) DNA demethylation and failed to significantly alter imprinted expression (data not shown). Although the treatment showed higher toxicity in NPCs, the limited DNA demethylation and preserved imprinted expression likely reflects slower cell division rates or additional imprint-preserving mechanisms in differentiated cells.

To identify additional epigenetic features associated with the allelic chromatin structural differences, we analyzed allelic histone modification datasets (**Figure 2G**). Active histone marks, including H3K27ac, H3K4me3, and H3K36me3, were preferentially enriched at unmethylated gDMRs, whereas the repressive histone mark H3K9me3 was associated with methylated gDMRs. H3K27me3 was also more prevalent at unmethylated gDMRs, consistent with its mutual exclusivity with DNA methylation. Together, these findings across multiple imprinted domains support a model in which methylation-sensitive CTCF binding at ICRs establishes chromatin insulation, and distinct histone modifications reinforce imprinted expression.

### Neural-specific chromatin structure at the *Meg3* locus

During our analysis of chromatin architecture in the cortex, we identified regions where differential allelic structures were not directly associated with CTCF binding. One notable example is the *Meg3-Dio3* imprinted domain, which contains a 210 kb stretch of chromatin corresponding to the *Meg3* lncRNA gene body that is devoid of long-range chromatin contacts specifically on the maternal allele (**Figure 3A**, grey highlight). The 5’ end of this region is anchored by a strong, maternal-specific CTCF peak, whereas no strong CTCF binding is detected at the 3’ end.

**Figure 3.**
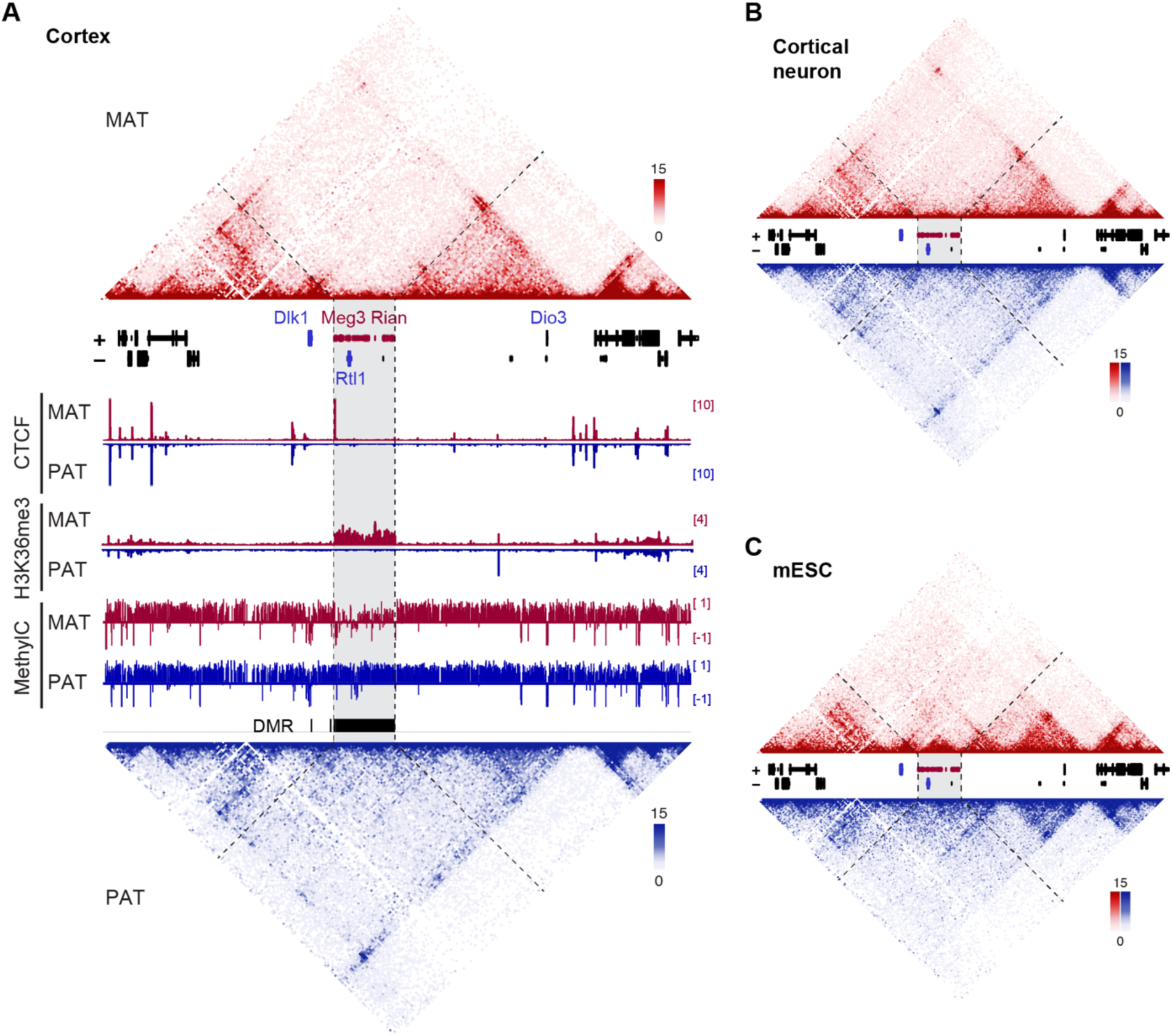
Neural-specific chromatin structure at the Meg3 domain. (**A**) Cortex capture Hi-C contact matrices, CTCF CUT&Tag, and MethylC tracks at the *Dlk1*-*Meg3* imprinted domain. (**B**) Primary cortical neuron and (**C**) mESC capture Hi-C contact matrices at the *Dlk1*-*Meg3* imprinted domain. Maternal tracks (red), paternal tracks (blue).

To investigate epigenetic features that might underlie this structural domain, we examined histone and DNA methylation landscapes. Although several imprinted lncRNAs, such as *Airn* and *Kcnq1ot1,* lead to deposition of H3K27me3 (Schertzer et al., 2019), the chromatin insulation pattern at the *Meg3* locus did not coincide with H3K27me3 enrichment. Instead, we found that maternal-specific enrichment of H3K36me3 mark correlated well with this region (**Figure 3A**, H3K36me3 track) along with an extended, heterogenous DMR exhibiting high DNA methylation on the inactive paternal allele and partially reduced methylation on the active maternal allele (**Figure 3A**, MethylC track).

A similar maternal-specific absence of long-range contacts was observed in primary cortical neurons (**Figure 3B**), whereas a more defined TAD-like structure was observed in mESCs (**Figure 3C**) and mESC-derived cardiomyocytes (Farhadova et al., 2024; Llères et al., 2019). These findings suggest that parental allele-specific chromatin structures can also arise from transcription-associated features, such as active transcription elongation and chromatin marks, in addition to classical CTCF-mediated loop formation.

### Allelic chromatin organization correlates with transcriptional activity and regulatory landscape at imprinted genes

Having established that imprinted domains exhibit distinct allele-specific chromatin architectures, we next investigated how these structural differences relate to imprinted gene expression. To examine the functional consequences of allelic chromatin organization, we asked whether imprinted genes show allelic differences in their contacts with distal regulatory elements. Because transcriptionally active promoters tend to exhibit higher interaction frequencies, or “hubness”, with regulatory elements, we quantified the frequency of distal chromatin contacts anchored at the transcription start sites (TSSs) of imprinted genes using our cortex capture Hi-C data (**Figure 4A**). Active and inactive alleles were defined based on allele-specific RNA-seq from hybrid primary cortical neurons (≥10 normalized reads across samples; **Extended Figure 3A**). As a control, we examined all biallelic (non-imprinted) genes located within the capture regions.

**Figure 4.**
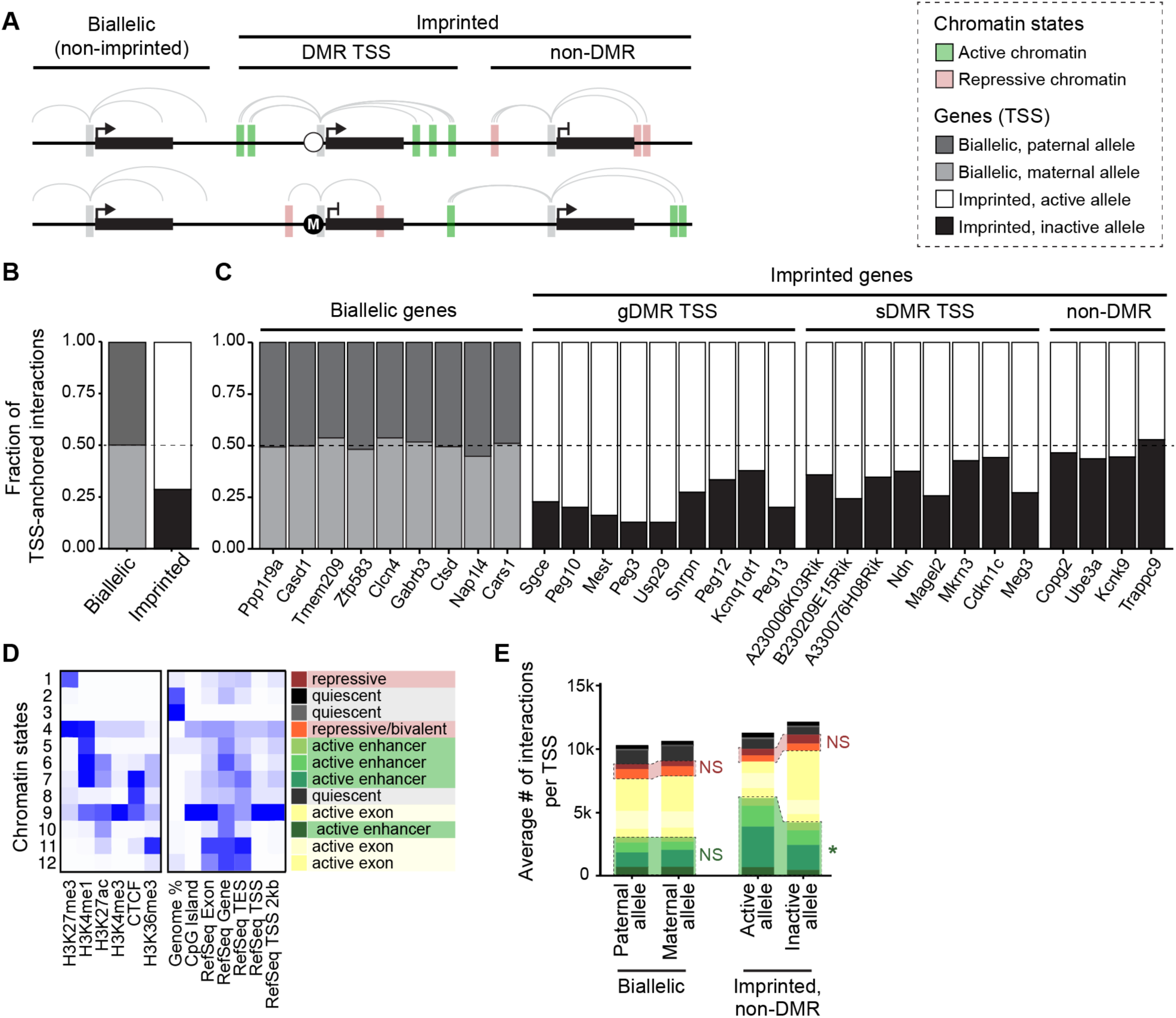
Imprinted gene promoters exhibit allelic differences in distal chromatin contacts. (**A**) Schematic of contact frequencies (arcs) from transcription start sites (TSS) at biallelic genes or imprinted genes with or without an overlapping DMR (plotted in B-C). Chromatin states at distal contact points illustrated as colored boxes (plotted in D-E). (**B**) Fraction of total TSS-anchored contacts from each allele of all biallelic or imprinted genes in the capture region. (**C**) Fraction of TSS-anchored contacts from each allele of individual biallelic or imprinted genes. Only a representative subset of biallelic genes is shown for illustrative purposes. (**D**) Chromatin states from ChromHMM and corresponding chromatin marks. Each state was assigned as quiescent, repressive, active enhancer, or actively transcribed exon. (**E**) Average number of TSS-anchored contacts per TSS, colored by chromatin state of distal contact point. Wilcoxon test was performed to compare the average contact frequencies between the two alleles for distal contacts marked by repressive or active chromatin states. n = 4 replicates of cortex capture Hi-C; NS, not significant; *p<0.05.

Biallelic genes exhibited similar frequencies of TSS-anchored distal contacts from each parental allele. In contrast, imprinted genes showed a marked imbalance: the active allele engaged in more distal chromatin contacts than the inactive allele (**Figure 4B**). When analyzed individually, genes whose TSSs overlapped gDMRs or sDMRs exhibited the strongest allele-specific differences in contact frequency compared with biallelic genes (**Figure 4C**). By contrast, imprinted genes lacking promoter-associated methylation (‘non-DMR’ genes) displayed similar contact frequencies from both alleles. Consistent observations were made from capture Hi-C in primary cortical neurons (**Extended Figure 3B,C**).

To better understand how non-DMR imprinted genes achieve allele-specific expression despite comparable contact frequencies, we next examined whether the regulatory potential differs between maternal and paternal contacts. Specifically, we assessed whether the chromatin state at the distal ends of TSS-anchored contacts differed between the two parental alleles (**Figure 4A**, colored boxes). To classify chromatin state by epigenetic landscape, we performed ChromHMM analysis (Ernst & Kellis, 2017) using publicly available ChIP-seq data together with our allelic CUT&Tag datasets, all from neuronal cells or tissue. The input features included allele-specific CTCF, H3K36me3, H3K27me3, H3K27ac and H3K4me3 profiles, as well as non-allelic H3K4me1 data. Chromatin states were inferred according to established conventions and grouped into four categories: “active enhancer,” “actively transcribing gene bodies,” “repressive,” and “quiescent” (**Figure 4D**).

We then quantified the average frequency of chromatin interactions from each TSS to ChromHMM-defined chromatin states. Biallelic genes exhibited minimal allelic differences in contact types, whereas the active alleles of the non-DMR imprinted genes preferentially interacted with “active enhancer” chromatin regions compared to inactive alleles in both cortex and primary cortical neurons (**Figure 4E, Extended Figure 3D**). Together, these results indicate that allele-specific gene expression is associated with high interaction frequency (“hubness”) and, in some cases, modest biases in the epigenetic states of distal interacting regions.

### CRISPRi screening uncovers functional enhancers at imprinted domains

Our Capture Hi-C analysis provided detailed maps of chromatin interactions at 5 kb resolution. However, because regulatory elements are often smaller than this scale, we next refined potential cis-regulatory regions within imprinted domains using chromatin accessibility profiling. ATAC-seq performed on hybrid mouse cortex identified approximately 78,000 open chromatin peaks genome-wide. Leveraging parental allele-specific SNPs, we examined whether any accessible regions exhibited allele-specific accessibility indicative of parent-specific regulatory activity. Around 100 ATAC peaks were exclusive to one parental allele, most of which overlapped known imprinted gene promoters or DMRs (**Extended Figure 4A**). Only a few exceptions were observed, including four intergenic peaks not associated with known imprinted domains and several allele-specific peaks within repetitive regions at the *Snrpn* locus. These findings suggest that allele-specific cis-regulatory elements outside of promoters and DMRs are rare, at least at the resolution and sensitivity of the current approach.

To identify functional cis-regulatory elements involved in imprinting, we next performed a CRISPR interference (CRISPRi) screen. We focused on the *Mest*-*Copg2* imprinted domain to investigate the regulatory basis of its allele-specific expression. Within this domain, *Mest* is expressed exclusively from the paternal allele, whereas *Copg2* shows a maternal expression bias specifically in the brain despite lacking a promoter-associated DMR. However, the potential contribution of cis-regulatory elements in this imprinted domain remains unknown.

We designed guide RNAs (gRNAs) targeting 25 accessible chromatin peaks within the *Mest*-*Copg2* domain (**Figure 5A, Supplementary Table 1**). Each peak was individually targeted with lentivirally delivered gRNAs in hybrid NPCs established from dCas9-KRAB x CAST F1 embryos. Upon neuron differentiation, transcript levels of *Mest* and *Copg2* were quantified by RT-qPCR (**Figure 5B**). Among the 25 candidate regions, inhibition of one element (hereafter referred to as E105) produced a significant reduction in *Mest* expression. Although SYBR-based RT-qPCR detected no change in total *Copg2* levels, allele-specific digital droplet PCR (ddPCR) demonstrated a significant reduction in the maternal-to-paternal *Copg2* expression ratio upon E105 inhibition (**Figure 5C**). This regulatory effect could not be explained by linear proximity, as E105 was not the closest element to either promoter. The E105 region also exhibits histone modifications characteristic of active enhancers, specifically H3K4me1 and H3K27ac, in the mouse brain (**Figure 5D**), supporting its enhancer identity.

**Figure 5.**
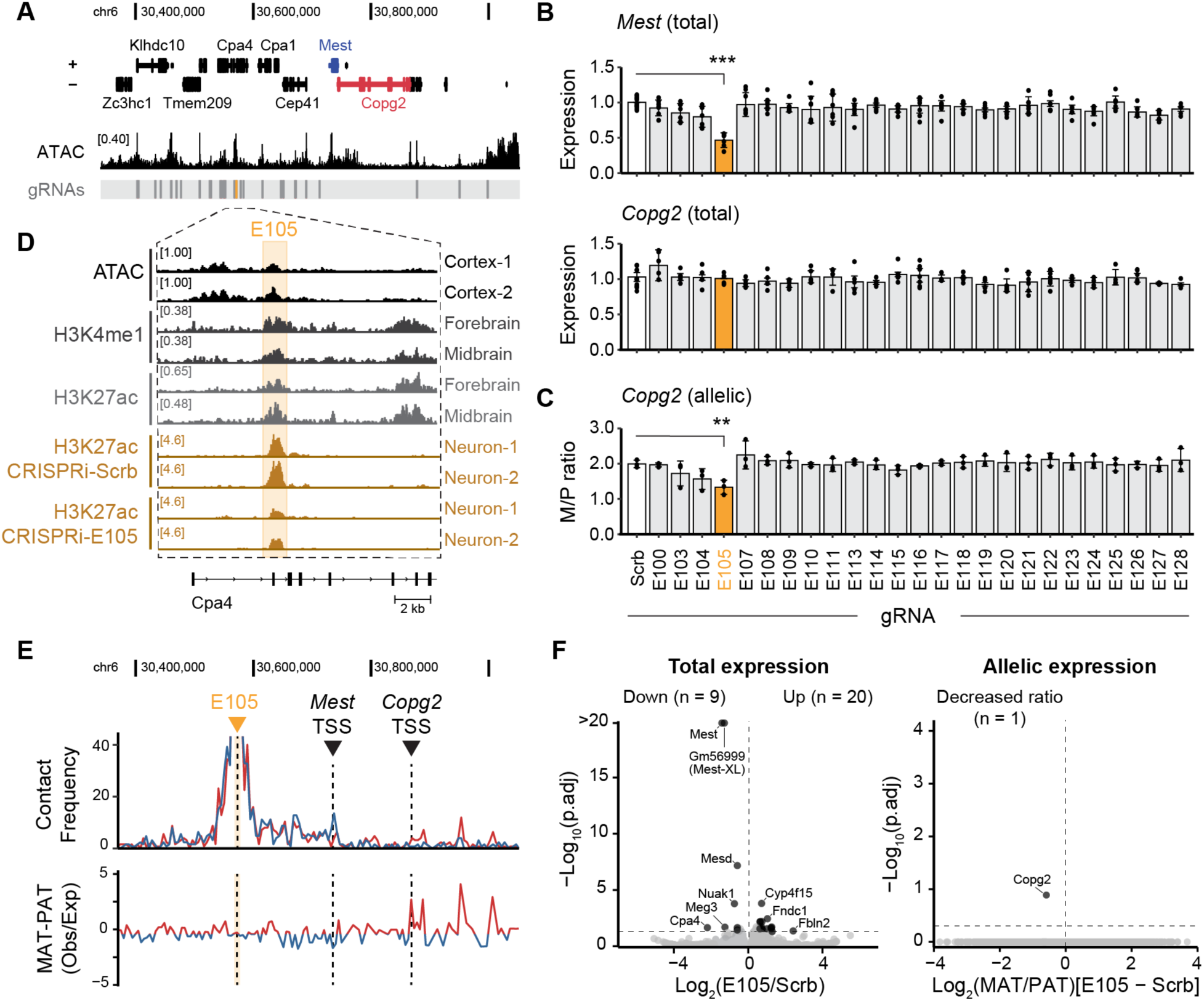
CRISPRi screening uncovers functional enhancers at imprinted domains. (**A**) ATAC-seq track and position of CRISPRi gRNA target sites in the *Mest*-*Copg2* domain. (**B**) Relative *Mest* or *Copg2* expression measured by qRT-PCR in differentiated NPCs upon CRISPRi for each target ATAC peak. Scramble (Scrb) gRNA is used for normalization. Bars represent the average of biological/independent experiments. All technical replicates shown in dots. Dunnett’s test *** < 0.001. (**C**) Ratio of allelic *Copg2* expression quantified by ddPCR using SNP-specific TaqMan probes upon CRISPRi for each target ATAC peak. Represented as maternal cDNA counts relative to paternal counts. Biological/independent replicates shown in dots. Dunnett’s test ** < 0.01. (**D**) Magnified view of E105 enhancer region and corresponding ENCODE H3K4me1 and H3K27ac histone modification tracks in mouse E15.5 forebrain and midbrain, as well as H3K27ac tracks in Scrb or E105 targeting gRNA transduced differentiated NPCs. (**E**) *Top:* Virtual 4C plot generated using capture Hi-C data anchored at E105 enhancer, showing the parental allele-biased contact frequencies. red = maternal contacts and blue = paternal contacts. *Bottom:* Distance normalized contact frequency anchored at E105 enhancer, red = maternally biased contacts and blue = paternally biased contacts. (**F**) *Left:* Volcano plot showing genes down-or up-regulated upon transduction of E105 targeting gRNAs in dCas9-KRAB NPCs followed by differentiation. *Right:* Volcano plot showing genes whose maternal to paternal ratio is differentially regulated upon transduction of E105 targeting gRNAs in dCas9-KRAB NPCs followed by differentiation. padj is treatment x allele interaction. n = 2 replicates per group.

Transcriptome-wide analysis confirmed that E105 functions as a highly specific regulatory element, with its inhibition selectively reducing *Mest* and altering the *Copg2* allelic bias while having minimal effects on global transcription (**Figure 5F**). CRISPRi of E105 resulted in a significant, localized loss of H3K27ac at the enhancer site (p_adj_ = 1.7×10^-13^; **Figure 5D**) without altering H3K27ac at the *Mest* or *Copg2* TSSs (p_adj_ > 0.95; **Extended Figure 4B-D**). Furthermore, deletion of E105 using dual gRNAs and catalytically active Cas9 recapitulated the reduction of *Mest* and altered *Copg2* allelic bias (**Extended Figure 4E**), validating the enhancer function of E105 in regulating the *Mest*-*Copg2* domain. To test the in vivo relevance of E105, we administered AAV constructs expressing GFP and E105-targeting gRNA constructs into adult dCas9-KRAB brain via retro-orbital injection. Approximately 45% of nuclei in the brain were GFP positive (**Extended Figure 4F**). RT-qPCR analysis using cortices from mice injected with the E105-targeting AAV showed a 20% reduction in *Mest* expression relative to scrambled controls (**Extended Figure 4G**).

Because *Mest* and *Copg2* display opposite parental expression patterns and our capture Hi-C maps revealed pronounced allele-specific domain organization, we next examined how these structural asymmetries might facilitate E105-mediated regulation of both genes. Visualization of chromatin contacts using virtual 4C plots revealed that E105 engages in paternally biased interactions with the *Mest* TSS and maternally biased interactions with the *Copg2* TSS (**Figure 5E**), consistent with enhancer-promoter communication through allele-specific chromatin loops. Collectively, these findings demonstrate that E105 functions as a long-range, allele-biased enhancer that coordinates *Mest* and *Copg2* expression within the same imprinted domain.

### *Copg2* is regulated by both a distal enhancer and *Mest*-XL lncRNA

A previous study demonstrated that tissue-specific *Copg2* imprinting is influenced by transcriptional interference from a 3’UTR-extended *Mest* isoform, *Mest-XL*, which is transcribed on the paternal allele antisense to *Copg2* in neural tissue (MacIsaac et al., 2012) (**Figure 6A**). To test whether *Mest*-*XL* functions as a nuclear lncRNA capable of modulating allele-specific regulation, we performed nuclear fractionation followed by RT-qPCR. *Mest-XL* was highly enriched in the nuclear fraction, whereas total *Mest*, representing both *Mest* and *Mest-XL* isoforms, showed only modest nuclear enrichment (**Figure 6B**). These results support a preferential nuclear localization and a potential non-coding role for *Mest-XL*.

**Figure 6.**
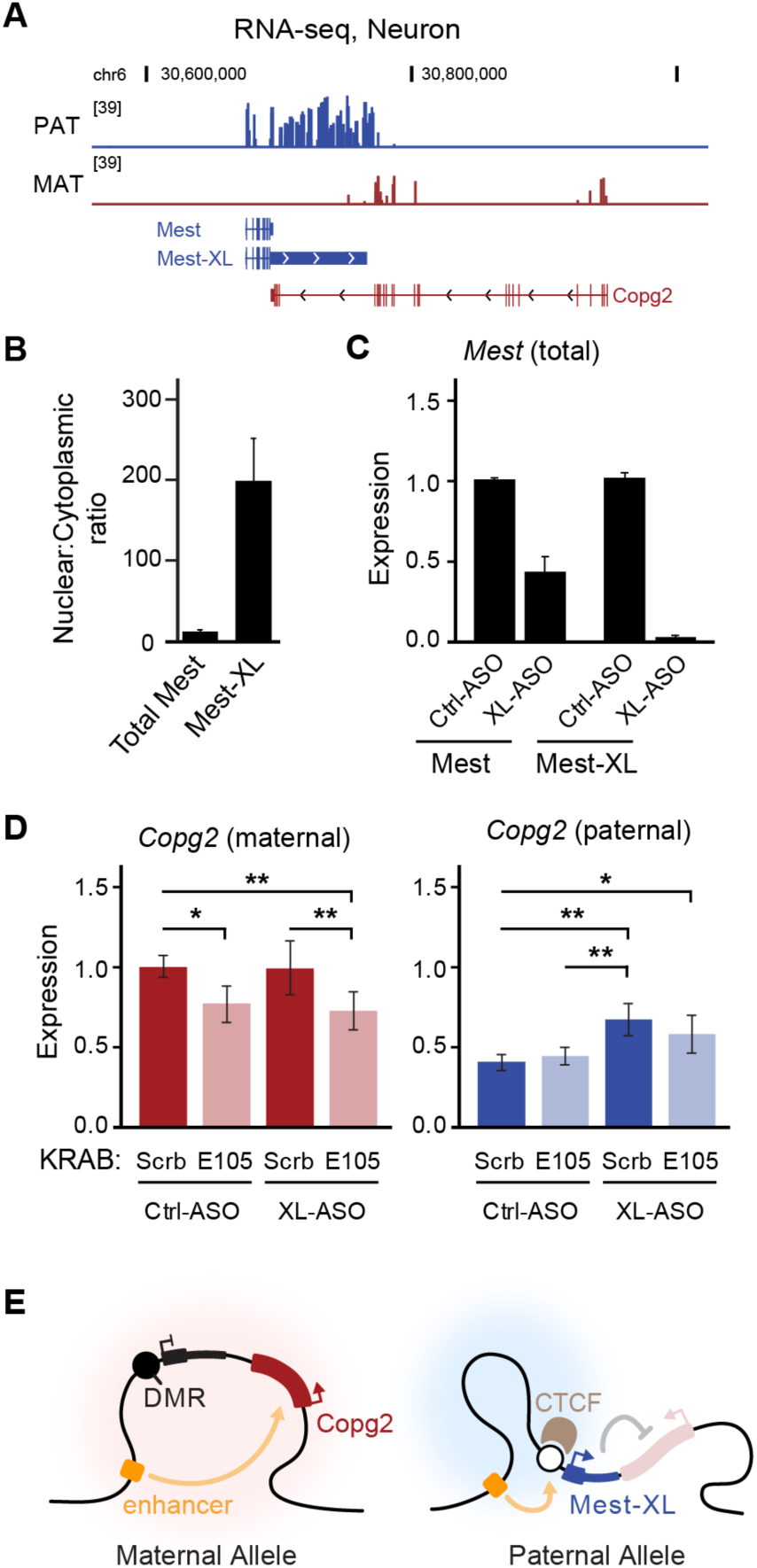
*Copg2* is regulated by both a distal enhancer and *Mest*-XL lncRNA. (**A**) RNA-seq tracks (log-scale axis) from primary cortical neurons showing the allelic expression of *Mest* and *Copg2*. (**B**) qRT-PCR result showing *Mest*-XL enrichment in the nuclear relative to cytoplasmic fraction. Nuclear fraction normalized to *Meg3* expression and cytoplasmic fraction normalized to *Psmd4* expression (n = 3). (**C**) Relative *Mest* expression measured by qRT-PCR upon free uptake of *Mest*-XL targeting ASO in NPCs followed by differentiation. (**D**) Allele-specific *Copg2* expression in differentiated NPCs quantified by ddPCR using allele-specific TaqMan probes. Samples are transduced with either Scrb vs E105 targeting gRNAs in combination with either Ctrl or *Mest*-XL targeting ASO. Maternal *Copg2* expression shown in red colors and paternal *Copg2* expression shown in blue colors. Tukey’s HSD-test * < 0.05, ** < 0.01. (**E**) Illustration of the *Mest*-*Copg2* regulatory mechanisms.

We next sought to determine if the effects of E105 enhancer on *Copg2* allelic bias were direct — via long-range contacts — or indirect through changes in *Mest-XL* transcription. Because inhibition of E105 reduced the maternal-to-paternal *Copg2* expression ratio, we asked whether this reflected decreased maternal expression or increased paternal expression. Absolute allele-specific quantification by ddPCR showed that E105 inhibition significantly decreased maternal *Copg2* transcript counts, accompanied by a relative increase in paternal transcripts (**Extended Figure 5A**). RNA-seq analysis yielded consistent results, with DESeq2-normalized counts indicating a maternal-specific reduction and paternal-specific gain in *Copg2* expression following E105 inhibition (**Extended Figure 5B**). Together, these data suggest that E105 directly activates the maternal *Copg2* allele, while repression of the paternal allele may involve *Mest-XL* transcription.

To disentangle the contributions of *Mest-XL* transcription and E105 enhancer activity, we selectively inhibited *Mest-XL* using antisense oligonucleotides (ASOs), which interfere with nascent transcription by inducing premature termination (Lee & Mendell, 2020). ASO treatment strongly down-regulated *Mest-XL* (>93%), resulting in ∼50% reduction in total *Mest* RNA (**Figure 6C**). Allele-specific ddPCR showed that *Mest-XL* knockdown selectively increased *Copg2* from the paternal allele, with no effect on the maternal allele (**Figure 6D**). In contrast, E105 inhibition by CRISPRi selectively reduced maternal *Copg2* expression, which was preserved both in the absence and presence of ASO treatment (**Figure 6D**). These findings clearly distinguish the contributions of the two regulatory components: E105 acts as a long-range enhancer that activates the maternal *Copg2* allele through allele-biased chromatin contacts. On the paternal allele, E105 instead engages the *Mest*/*Mest-XL* promoter region, contributing to *Mest/Mest-XL* transcriptional output. *Mest-XL* then represses paternal *Copg2* through transcriptional interference. Thus maternal activation by E105 and paternal repression by *Mest-XL* operate on opposite alleles and mutually reinforce each other to establish tissue-specific *Copg2* imprinting (**Figure 6E**).

## Discussion

Our study reveals that robust, allele-biased chromatin conformations are pervasive across multiple imprinted domains in the mouse brain, establishing allele-specific 3D genome organization as a general feature of imprinting regulation. Consistent with classical models, unmethylated ICRs serve as strong chromatin boundaries marked by allele-specific CTCF binding, while methylated alleles lack insulation and exhibit more continuous interaction profiles. Loss of DNA methylation at ICRs is sufficient to induce biallelic CTCF binding, disrupt allele-specific chromatin architecture, and impair imprinted expression, underscoring the direct role of methylation-sensitive insulation in establishing parental-specific chromatin domains. These results reinforce the long-standing paradigm that ICRs are master regulators of imprinting but also highlight how allele-specific chromatin domains contribute to regulatory specificity across multiple imprinted loci.

Beyond ICRs, we find that allele-specific chromatin organization also modulates distal regulatory engagement, extending the imprinting regulatory model to sub-TAD scales. Active alleles of imprinted genes engage in more frequent distal contacts than inactive alleles, providing a structural correlate of transcriptional “hubness.” Notably, imprinted genes lacking promoter-associated DMRs show minimal differences in contact frequency but display allele-biased differences in the regulatory potential of contacted chromatin, as revealed by ChromHMM-defined epigenetic states. These findings indicate that imprinted expression can arise not only from differential promoter interaction frequency but also from subtle, allele-biased interactions with distal regulatory elements. Importantly our analysis across multiple imprinted domains enables systematic comparison of allele-specific chromatin interactions and their functional relevance, strengthening the connection between 3D genome organization and imprinted regulation.

Through a CRISPRi screen of accessible elements at the *Mest–Copg2* domain, we identify a distal enhancer, E105, that exerts highly specific regulatory effects on *Mest* and *Copg2*. E105 engages in allelic long-range interactions with both promoters, showing paternally biased contacts with *Mest* and maternally biased contacts with *Copg2*, revealing that a single enhancer can be differentially employed on the two parental chromosomes. These asymmetric interactions, together with enhancer-associated histone marks and loss-of-function perturbations, support a model in which E105 functions as a shared enhancer whose activity is partitioned by allele-specific chromatin architecture.

This architectural logic is similar to the well-established paradigm at the *H19*–*Igf2* locus, where a methylation-sensitive ICR sets the boundary that governs allele-specific enhancer access. However, our data also reveal a key distinction: whereas the *H19*–*Igf2* ICR functions exclusively as an insulator, the *Mest*–*Copg2* ICR appears to serve a dual role. In addition to restricting E105 from the paternal *Copg2* promoter, it also anchors E105 contacts on the paternal chromosome to support *Mest*/*Mest*-XL transcription. This dual function likely reflects the promoter-overlapping position of the *Mest* ICR and suggests that certain ICRs can participate directly in allele-specific regulatory looping rather than acting solely as boundaries.

Interestingly, E105 inhibition altered *Copg2* allelic ratios without changing total transcript levels, raising the possibility that interallelic compensation or feedback maintains total gene output. Understanding how alleles communicate to preserve overall dosage represents an important open question for imprinting biology.

Importantly, our findings integrate the function of E105 with the known paternal repression mediated by *Mest-XL*, a neural-specific antisense transcript previously shown to influence *Copg2* imprinting via transcriptional interference (MacIsaac et al., 2012). By combining enhancer inhibition and *Mest-XL* knockdown, we demonstrate that these two mechanisms act on opposite alleles: E105 promotes maternal activation, whereas *Mest-XL* represses the paternal allele. Thus, *Copg2* imprinting emerges from the combined action of enhancer-driven activation and lncRNA-driven repression, coordinated through an allele-specific chromatin environment. This dual mechanism provides a compelling explanation for how *Copg2* achieves tissue-specific maternal expression despite lacking a promoter DMR.

Together, our results broaden the mechanistic landscape of imprinting beyond canonical ICR-driven insulation. Distal enhancers, allele-specific chromatin structures, and lncRNA-mediated transcriptional interference jointly create a multilayered regulatory architecture that fine-tunes imprinted expression in a tissue-specific manner. These principles likely extend to other imprinted loci and non-imprinted genes with allele-biased expression, particularly those embedded in complex chromatin neighborhoods.

Future work will be needed to determine the generality of enhancer–lncRNA cooperation at other imprinted domains, to delineate how allele-specific chromatin loops are dynamically regulated across development, and to define the interplay between transcriptional activity and chromatin topology in reinforcing parent-of-origin–specific expression patterns. More broadly, our findings underscore the importance of integrating 3D genome architecture with functional genomics to fully understand the epigenetic logic of monoallelic gene regulation.

## Methods

### Standard mouse housing

All mice were maintained in accordance with the “Guide for the Care and Use of Laboratory Animals” (National Research Council, 8^th^ edition, 2011) and all animal protocols were approved by the Harvard University FAS IACUC. Mice were housed at 22 +/- 1°C with 30-70% humidity and a 14L:10D cycle.

### Primary cell culture

For NPC culture, the dorsal region of mouse cortices from E14 embryos were carefully collected into HEPES buffer saline solution. Tissues were dissociated with 0.25% Trypsin/EDTA at 37°C for 5 min and assisted by mechanical trituration by pipetting. Cell suspensions were filtered through a 40 μm cell strainer, then plated onto sterile bacterial petri dish and incubated for 4-5 days to induce neurosphere formation. Media change was performed every day during this period. For adherent NPC culture, neurospheres were dissociated with 0.25% Trypsin/EDTA at 37°C for 5 min. Dissociation was assisted by mechanical trituration with pipette in the presence of defined trypsin inhibitor. Cells were maintained onto Geltrex-coated tissue culture dishes for 3-4 passages. NPC were cryopreserved until use.

For primary cortical neuron culture, cortices from E16 embryos were carefully collected into cortical neuron plating media (Neurobasal supplemented with 2% B27, 2% FBS and 1X glutamine). Tissues were mechanically triturated into the plating media by pipetting up and down with 1000 μL micropipette tips. Cell suspensions were filtered through a 40 μm cell strainer. Dissociated cells were plated onto Poly-D-lysine coated plates with plating media. The next day, media was replaced with cortical neuron maintenance media (Neurobasal supplemented with 2% B27 and 1X glutamine) and maintained at 37°C with 5% CO_2_ until use.

### mESC culture

For routine maintenance of mouse ESCs, cells were grown on 0.2% gelatin coated plate in ESC media (DMEM supplemented with 10 mM HEPES, 0.11 mM β-mercaptoethanol, 1X NEAA, 2 mM L-glutamine, 15% FBS, and 1000 U/mL LIF). For passage, cells were washed with HEPES-buffered saline, dissociated using 0.25% Trypsin/EDTA, and appropriate dilutions were made to plate as routinely. For drug treatment, 5 μM GSK-3484862 was added to treatment groups and volume matched DMSO was added to controls. Cells were maintained at 37°C with 5% CO_2_.

### Capture Hi-C library preparation

For Hi-C library preparation, Arima Hi-C kit was used following provider’s instruction. Dissected mouse cortices (two BL6xCAST and two CASTxBL6; 7-8 weeks-old) were snap frozen in liquid nitrogen and stored at −80°C. Frozen tissues were pulverized using Cell crusher in liquid nitrogen. Adherent cells were washed with PBS two times and scraped into 5 mL PBS. Samples resuspended in 5 mL PBS and crosslinked in 2% formaldehyde by adding 500 μL of TC buffer (HEPES 50 mM, NaCl 100 mM, EDTA 1 mM, 0.5 mM EGTA, and 22% formaldehyde) and incubating at room temperature for 20 min. Reaction was quenched by adding 289 μL of strop solution I and incubating at RT for 5 min. Crosslinked tissue powder was then centrifuged at 2,000 xg at RT for 15 min and supernatant was discarded. Pellet was stored at −80°C until use. Samples were resuspended in 20 μL lysis buffer, lysed at 4°C for 30 min, conditioned using SDS solution at 62°C for 10 min, and quenched by incubating with 20 μL of stop solution II at 37°C for 15 min. Digestion of extracted chromatin was performed using Arima enzymes by incubating at 37°C 60 min, 65°C 20 min, and 25°C 10 min. Digested chromatin was subject to biotin end repair at room temperature for 45 min, followed by proximity ligation at room temperature for 15 min. Crosslinking reversal was performed using proteinase K and incubating at 55°C 30 min, 68°C overnight, and 25°C 10 min. Proximity-ligated DNA was purified using AMPure XP beads. Proximity-ligated DNA was brought to 100 μL in elution buffer and fragmented using Covaris S220 (140 W, 10% duty factor, 200 cycles per burst, 55 sec) to obtain a target fragment of 400 bp. Double sided size selection was performed using 0.6X and 0.25X AMPure XP beads. Biotin enrichment was performed using 300-400 ng of size-selected DNA and 12.5 μL T1 Streptavidin beads in 200 μL binding buffer reaction. Two washes were performed in 200 μL wash buffer at 55°C for 2 min each. Final wash was performed in 100 μL of elution buffer. Washed beads were resuspended in 50 μL of water. On beads end repair was performed by incubating end repair mix at 20°C 15 min 72°C 15 min, followed by adaptor ligation at 20°C 30 min. Samples were cleaned up by magnetizing beads and washing in 200 μL wash buffer at 55°C 2 min and in 100 μL elution buffer. Washed beads were resuspended in 34 μL of water and immediately used for library amplification using Herculase II Fusion polymerase (98°C 2 min / 10X 98°C 30 sec 60°C 30 sec 72°C 1 min / 72°C 5 min). Different index primer combinations were used for each sample. Amplified library was purified using 1X AMPure XP beads. An aliquot of 1.5 μg PCR product in 30 μL elution buffer was pre-cleared of biotin using 50 μL T1 Streptavidin beads in 230 μL reaction. Pre-cleared Hi-C library was resuspended in 14 μL water. Approximately 1 μg of Hi-C library was incubated with custom probes (Agilent Technologies) in thermocycler (95°C 5 min, 65°C 10 min / 60X 65°C 1 min 37°C 3 sec) and purified using 50 μL T1 Streptavidin beads. Vigorous washes were performed six times at 70°C 5 min each. Capture enriched library is amplified using Herculase II polymerase (98°C 2 min / 13X 98°C 30 sec 60°C 30 sec 72°C 1 min / 72°C 5min). Final library purification was performed using 1X AMPure XP beads and resuspended in 25 μL water. Samples were pooled equimolar. Illumina PE150 sequencing was performed (Novogene).

### Capture Hi-C analysis

Mouse genome GRCm39 was prepared using SNPsplit_genome_preparation script to mask all the CAST_EiJ single nucleotide polymorphism positions using mgp_REL2021_snps.vcf.gz from the Mouse Genomes Project (Keane et al., 2011; Krueger & Andrews, 2016). SNP-masked genome was digested at Arima restriction enzyme sequences using HiCUP_digester (Wingett et al., 2015). Capture Hi-C fastq files were mapped using HiCUP (v0.8.0). This wrapper identified mapped paired-end reads with valid Hi-C di-tags. Allele phasing was performed inputting ‘hicup.bam’ files to SNPsplit (v0.6.0). Read pairs in ‘hicup.G1_G1.bam’ or ‘hicup.G2_G2.bam’ files were converted to ‘prejuicer’ format using HiCUP hicup2juicer script (v0.9.0) with --usemid - -digest options. Using a custom script, reads were filtered for the capture regions with MAPQ score > 30 at both ends, and maternal and paternal read pair numbers were normalized by subsampling the deeper allele. Medium files were independently assessed to verify reproducible interactions. Otherwise, files were merged according to its parental origin (i.e., ‘G1_G1.medium’ files from BxC and ‘G2_G2.medium files from CxB samples were merged). For comparison between different cell/tissue data, capture region filtered reads were subsampled to equalize the depth (i.e. 158,618 read pairs in relation to **Figure 3**). Medium files were converted to hic format using Juicer pre tool (v1.22.01). Visualization of contact matrix was conducted in R (v. 4.4.1) using plotgardener (v1.10.2) (Kramer et al., 2022) or Juicebox (v. 1.11.08). Chromatin loop prediction was performed using Juicer HICCUPS using VC_SQRT normalization (Rao et al., 2014). Pileup analysis was performed using Coolpuppy (Flyamer et al., 2020). For chromatin contact frequency analysis, chromatin states were first classified using ChromHMM (Ernst & Kellis, 2017) with allele-specific CTCF, H3K36me3, H3K27me3, H3K27ac, and H3K4me3 ChIP-seq or CUT&Tag data and non-allelic H3K4me1 ChIP-seq data. Then, chromatin contacts corresponding to each state group were counted.

### CUT&Tag library preparation

Cells were gently washed with PBS and scraped into 1 ml PBS. An aliquot equivalent of 600,000 cells were used for each CUT&Tag sample. Cells were collected by centrifugation for 4 min at 300xg and supernatant was discarded. Cells were resuspended in lysis buffer (10 mM Tris pH 7.4, 10 mM NaCl, 0.5 mM spermidine, 1.5 mM MgCl, 0.1 mM EDTA, and 0.25% IGEPAL CA-630) and incubated for 5 min on ice. Lysis was assisted mechanically by gently pipetting up and down 10 times with 1000 μL tips. Successful lysis was confirmed by Typan blue staining and inspection under a light microscope. Nuclei were collected by centrifugation for 4 min at 800xg. Nuclei pellet was washed with CUT&Tag wash buffer and resuspended in 1.5 mL wash buffer. All the procedures were performed according to manufacturer’s guides. Concanavalin A beads were conditioned by washing with 1.6 mL 1X CUT&Tag binding buffer two times. Prepared beads were resuspended in binding buffer and slowly added to the nuclei suspension. For each sample, 20 μL of beads were used. Nuclei binding was performed for 10 min at room temperature in a rotator. Beads-bound nuclei were collected using a magnetic stand and clear supernatant was discarded. Beads were resuspended in 50 μL ice-cold CUT&Tag antibody buffer and desired amount of antibody was added (2 μL CTCF antibody Cell Signaling #2899, 1 μL H3K36me3 antibody Active Motif #61902, or 1 uL H3K27ac antibody Active Motif #39136). Suspension was gently mixed by vortexing. Primary antibody binding was performed overnight at 4°C. After a quick spin, beads were cleared on a magnetic stand and supernatant was discarded. Beads were resuspended in 100 μL of 1:100 diluted secondary antibody in CUT&Tag Dig-Wash buffer and incubated for 1h at room temp with rotation. After secondary antibody binding, three washes with 1 mL CUT&Tag Dig-Wash buffer were performed. Beads were resuspended in 1:100 diluted CUT&Tag transposomes in CUT&Tag Dig-300 buffer and incubated on an orbital rotator for 60 min at room temperature. After transposome binding, beads were washed three times with Dig-300 buffer. Tagmentation was induced by incubating beads in 125 μL of CUT&Tag tagmentation buffer for 1h at 37°C. After tagmentation, total DNA was extracted using columns. Purified DNA was PCR amplified using proper index primers. Then, the PCR reaction was purified using SPRI beads. Samples were pooled equimolar. Illumina PE150 sequencing was performed (Novogene).

### CUT&Tag analysis

Adapters sequences were automatically detected and trimmed using TrimGalore (v0.6.10). Reads were mapped to CAST SNP-masked GRCm39 genome using bowtie2 --local -X 2000 (v2.5.1). PCR duplicates were removed using Picard MarkDuplicates. Allele-specific reads were parsed using SNPsplit. Peak calling was performed using MACS2 (v2.2.7.1) or SICER2 callpeak inputing un-split bam files. Peaks from each bam file were merged and converted to saf format for its use in FeatureCounts (subread v2.0.6). Read counts were performed by featureCounts -p -- countReadPairs using SNP split bam files. Parental biased read enrichment was assessed in R using DEseq2. For visualization purposes, bam files were converted to bigwig files with CPM normalization (deeptools v3.5.2). Differential analysis was performed in R using DEseq2 (v1.40.2).

### ATAC-seq library preparation

ATAC-seq library was prepared using the ActiveMotif ATAC-seq kit. Frozen mouse brains were minced with razor blade on ice and quickly transferred to conical tubes containing cold PBS. Tissue was centrifuged at 500 xg for 5 min at 4 °C. PBS was discarded, and the pellet was resuspended in 1 mL ATAC lysis buffer. Lysis was assisted by dounce homogenizer with tight fitting pestle for 30 strokes. Samples were filtered through 40 μm strainer into a new conical tubes. Nuclei were counted using trypan blue, and 100,000 nuclei were transferred to a new tube. Nuclei were centrifuged at 500 xg for 5 min at 4°C, and supernatant was discarded. Nuclei pellets were subject to tagmentation at 37°C for 30 min with gentle shaking at 800 rpm. Immediately following the tagmentation reaction, total DNA was extracted using columns. Purified DNA was PCR amplified using proper index primers. Then, the PCR reaction was purified using SPRI beads. Samples were pooled equimolar. Illumina PE150 sequencing was performed (Novogene).

### ATAC-seq analysis

Adapters sequences were automatically detected and trimmed using TrimGalore. Reads were mapped to CAST SNP-masked GRCm39 genome using bowtie2 --very-sensitive -X 2000. PCR duplicates were removed using Picard MarkDuplicates. Allele-specific reads were parsed using SNPsplit. Peak calling was performed using MACS2 callpeak inputing un-split bam files. Peaks from each bam file were merged and converted to saf format for its use in FeatureCounts. Read counts were performed by featureCounts -p --countReadPairs using SNP split bam files. Parental biased read enrichment was assessed in R using DEseq2.

### RNA-seq analysis

RNA was extracted using Trizol (Thermo Fisher Scientifics) method. DNA was removed from the samples using Turbo DNase (Thermo Fisher Scientifics). Library preparation and sequencing was performed by Novogene. mESC RNA-seq upon DMSO or GSK-3484862 treatment was performed using poly(A) selection strategy. RNA-seq using cortical neurons or differentiated NPCs upon CRISPRi were performed using rRNA depletion strategy. Adapters sequences were automatically detected and trimmed using TrimGalore. Reads were mapped to CAST SNP-masked GRCm39 genome using STAR –sjdbGTFfile Mus_musculus.GRCm39.110.gtf -- sjdbOverhang 149 --alignEndsType EndToEnd (v2.7.0e). PCR duplicates were removed using Picard MarkDuplicates. Allele-specific reads were parsed using SNPsplit. Read counts were performed by featureCounts -p –countReadPairs -t exon -g gene_id -a Mus_musculus.GRCm39.110.gtf and using SNP split bam files. Parental biased read enrichment was assessed in R using DEseq2.

### gRNA cloning

Two to four gRNAs targeting select ATAC peaks were designed using CRISPick (Broad Institute). gRNAs were selected based on reduced overlap to increase the target range. gRNAs were cloned into 4X gRNA system (Kabadi et al., 2014) with some adaptations. The gRNA scaffold sequences in each of the gRNA plasmids (i.e. ph7SK, phH1, phU6, and pmU6) were replaced with the one from CROP-seq-opti plasmid to increase dCas9-KRAB residence time (Hill et al., 2018). gRNA flanking overhang sequences were also mutated to CACC to increase interchangeability of gRNAs. gRNAs were cloned into each plasmid using digestion and ligation method. Constructs were amplified using Q5 high fidelity PCR and were subject to HIFI assembly in order to clone all 4 constructs into a single lentiviral vector. The lentiviral vector pLV-GG-hUbC-dsRed was adapted to replace fluorescence marker with blasticidin resistance gene to increase compatibility with the dCas9-KRAB mouse system. Cloning of all four constructs were confirmed by colony PCR and whole plasmid sequencing or Sanger sequencing.

### Lentivirus production

For lentivirus generation, 293T (ATCC) cells were always maintained at 50-80% confluency in DMEM supplemented with 10% FBS. On the day of transfection, cells were plated onto 6 well plates at 80% confluency and incubated for 3-4 hours to allow cell adherence. 500 ng of pMD2G, 1000 ng of psPAX2, and 1500 ng of transfer plasmid were co-transfected using 9 μL PEI in OptiMEM. The next day, media was replaced with 1 mL of cortical neuron culture media. After 48 hours, supernatant was collected and centrifuged at 800xg for 4 min. Cleared supernatant were aliquoted and stored at −80°C to reduce freeze thaw cycles. All aliquots were used within 3 freeze thaw cycles. Lentiviral titration was performed using Lenti-X qRT-PCR Titration Kit (Takara Bio).

### Lentivirus transduction

NPCs were plated onto Geltrex coated plates at a density of 50,000 cells/mL (e.g. in 12 well plates 50,000 cells with 1 mL media, and in 6 well plates 100,000 cells in 2 mL media) and incubated for 3-4 hours to allow cell adherence. Transduction was performed at 7500 genomic particles per cell. The next day, NPC media was replenished to keep the multipotent state of the NPCs. After 48h of transduction, cells were replated onto new Geltrex coated plates with blasticidin-supplemented media. The next day, neuron differentiation was induced by replating NPCs onto PDL coated plates with cortical neuron media with blasticidin. The following day, cell culture media was optionally replaced to fresh cortical neuron media to enhance neuronal differentiation. On day 3 of differentiation, samples were collected into 200 μL of DNA/RNA shield solution (Zymo).

### AAV injection

pAAV-U6-sgRNA-CMV-GFP plasmid was used as backbone. CMV promoter was replaced by UbC promoter and 3x SV40 NLS were inserted at the N-terminus of GFP. Scrb gRNA was cloned downstream of hU6 promoter and 2 gRNAs targeting E105 were cloned under the control of hU6 and hH1 promoters. AAV.PHP.eB were packaged by Boston Children’s Hospital Viral Core. 5E11 AAV per adult animal (dCas9-KRAB; The Jackson Laboratory #030000) were injected retro-orbitally. After 3 - 3.5 weeks, mice were euthanized using isoflurane overdose and appropriate procedures were performed for each experiment.

### Microscopy

After isoflurane overdose, mice were perfused with PBS and 4% PFA. Brains were dissected out and fixed in 1% PFA overnight at 4°C. Fixed brains were washed in PBS and submerged in 30% sucrose overnight at 4°C. Brains were additionally incubated in a 1:1 mixture of OCT and 30% sucrose for 30 min to 6 hours. Brains were embedded in OCT and kept in – 80°C until use. Cyosections were prepared at 12-14 microns onto super plus frost slides. For AAV-derived GFP imaging, nuclei were stained using Hoeschst 33342 and mounted using ProLong Glass Antifade Mountant. Images were taken using Zeiss Axio Scan Z1 at 10X magnification. Total and GFP positive nuclei counting was performed in ImageJ using particle analysis tool.

### Antisense oligonucleotide treatment

Affinity plus gapmer ASO targeting the 3’ extension of *Mest-XL* (+T*+A*+C*A*T*T*T*G*A*G*T*C*T*+C*+C*+T) or scramble control ASO (+T*+C*+C*C*T*G*A*A* G*G*T*T*C*+C*+T*+C) were synthesized and purified by IDT. 250 μM stock ASO was diluted in NPC culture media to bring it to 12.5 μM final concentration and allowed free uptake by NPCs. Next day, NPC media was fed with half the original media volume. After 3 days of ASO treatment, NPC was replated onto PDL coated plates in primary cortical neuron maintenance media to allow differentiation. The next day, full media change was performed, supplemented with 250 μM ASO. Samples were collected on day 3 of differentiation unless otherwise stated.

### SyBR qRT-PCR

RNA was extracted using Trizol (Thermo Fisher Scientifics) method. DNA was removed from the samples using Turbo DNase (Thermo Fisher Scientifics). RNA was diluted at 5 ng/μL, and 2 μL of diluted RNA was used in one step RT-qPCR either using iTaq Universal One-Step RT-qPCR kit (BioRad) or Luna Univesal One-Step RT-qPCR kit (NEB). qPCR analysis was performed using ddCt method. *Psmd4* gene was used for normalization across samples. Primer sequences are listed in **Supplementary Table 2**.

### Allele-specific digital droplet PCR

RNA was extracted using Trizol (Thermo Fisher Scientifics) method. DNA was removed from the samples using Turbo DNase (Thermo Fisher Scientifics). 300-350 ng of total RNA was subject to cDNA synthesis using Maxima reverse transcriptase (Thermo Fisher Scientifics) and random hexamers. cDNA was diluted 1:5 in water. 5 μL of diluted cDNA was mixed with 1X TaqMan probes (Thermo Fisher Scientifics; SNP genotyping probes Assay ID AN49G3G – probe 1 CTCAGAAGACTTAAACAAG, probe 2 TCAGAAGACTTGAACAAG, forward amplification primer CTGCCATTCCTGAGTTTATGAACCT, and reverse amplification primer CGAAATACTCTGTCTCTGCTTCTGT) and 1X ddPCR Supermix for Probes (no dUTP) (BioRad 186-3023). *Ubc* gene was used for normalization across samples. Droplet generation was performed using QX200 droplet generator (BioRad). Droplets were carefully transferred to thermocycler and PCR was performed at 95°C 10 min / 40X 94°C 30 sec 60°C (with ramp rate +2°C /sec) 1 min / 98°C 10 min. Droplet count was performed within 1 hour of PCR using QX200 droplet reader (BioRad) with absolute quantification program.

### gDNA extraction

Cells were resuspended in 300 μL cell lysis buffer (0.1 M Tris pH 8, 0.2 M NaCl, 5 mM EDTA, 0.1% SDS, and 10 units/mL Proteinase K) and incubated at 50°C 1h to overnight. gDNA was precipitated with 600 μL of 100% isopropanol and incubated at −20°C for 2 hours before centrifuging at 12,000xg for 15 minutes. The supernatant was carefully discarded, and the remaining pellet was washed with 500 μL of 70% ethanol. Samples were again centrifuged at 12,000xg for 15 minutes, supernatant was discarded, and the pellet was resuspended in 50 μL of water.

### Bisulfite treatment and nanopore methylation analysis

Bisulfite treatment was performed using EpiTect Fast DNA Bisulfite kit (Qiagen 59824) following manufacturer’s instruction. Bisulfite converted DNA was eluted in 15 μL of water. Target fragment was PCR amplified using EpiMark Hot Start Taq (NEB). 1.5 μL of eluted DNA was used for each reaction. PCR was performed at 95°C 30 sec / 45X 95°C 30 sec 45°C 30 sec 68°C 30 sec / 68°C 5 min using bisulfite conversion-aware primers (**Supplementary Table 2**). PCR product was gel purified and pooled for sequencing at Plasmidsaurus using Premium PCR sequencing service. Fastq files were mapped to Cast SNP masked and CT converted mouse GRCm39 genome using minimap2 (v2.24). Reads were split according to SNPs to phase reads originated from maternal vs paternal allele. For each allele, CpG methylation percentage was calculated.

### Sanger sequencing

PCR was performed using Q5 High-Fidelity 2X Master Mix (NEB). Samples were run on 1% agarose gel and visualized using Gel Loading Dye, Purple (6X) (NEB). Fluorescent bands were cut from the gel and extracted using NucleoSpin Gel and PCR Clean-up kit (Macherey-Nagel) before sanger sequencing. Sanger sequencing was performed by Eton Biosciences.

### Statistics and reproducibility

Statistical analyses, including Wilcoxon, Dunnett’s, Tukey’s HSD, and t-test, were performed using R 4.4.1, with significance assumed if p < 0.05. The number of replicates is indicated in the figure legends.

## Acknowledgments

We thank all the members of the Whipple lab, Sofia Battaglia (Bernstein lab), Tricia Horvath (Dulac lab), Brandon Logeman (Dulac lab), Jenny Chen (Eddy lab), Patti Murphy (McKinley lab), and Matt Pezone (Mooney lab) for providing feedback, technical assistance, and/or experimental advice for this project. We also thank the Boston Children’s Hospital Viral Core (supported by NIH5P30EY012196) and The Bauer Core Facility at Harvard University for technical support and access to instrumentation. This work was supported by NIGMS R35GM146921 (A.J.W), George W. Merck Fellowship Fund (A.J.W), Rita Allen Foundation (A.J.W) and NIGMS F32GM156080 (B.B).

## Author Contributions

B.B. and A.J.W. conceptualized the project, B.B., K.G., and D.L. performed investigation and formal analysis, B.B. and K.G. wrote the initial manuscript, A.J.W. performed review and editing of the text, A.J.W. supervised and acquired funding for the work.

## Data availability

The datasets supporting the conclusions of this article have been deposited in the Gene Expression Omnibus under accession identifiers GSE312071, GSE312069, GSE312070, GSE312072, and GSE312073.

---

**Extended Figure 1.**
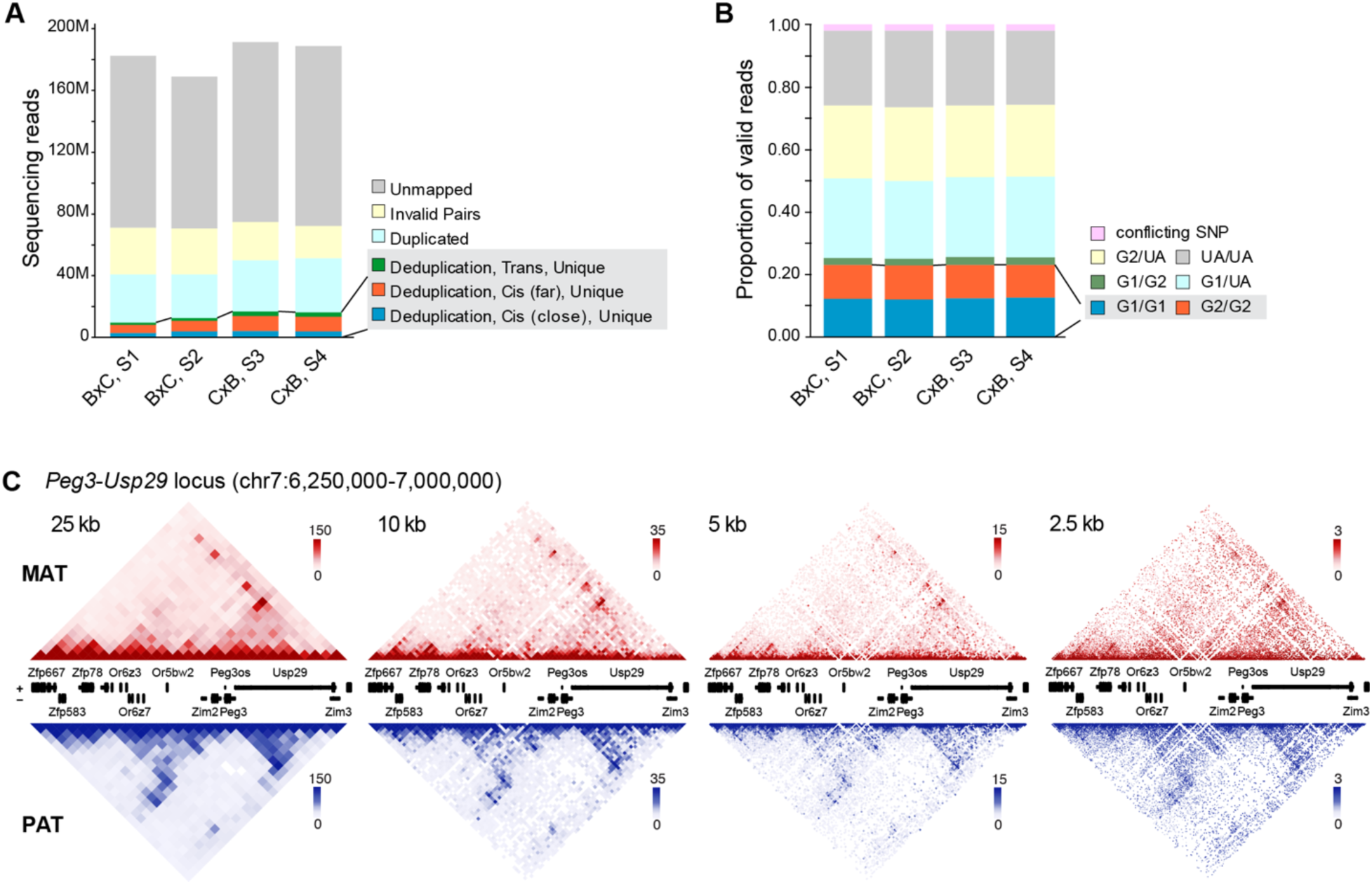
Imprinted domains exhibit pervasive parental allele-specific chromatin architectures. (**A**) Number of paired sequencing reads generated by Capture Hi-C from mouse cortex. Successfully deduplicated and ligated hybrid reads (highlighted in gray) were used for parental allele phasing. (**B**) Deduplicated hybrid reads were phased according to parental SNPs using SNPsplit. Intrachromosomal hybrid reads containing parental-specific SNPs (i.e. G1-G1 and G2-G2) were used for generating Hi-C contract matrices. G1, genome 1 (BL6); G2, genome 2 (CAST); UA; unassigned. (**C**) Example contact matrices of the *Peg3*-*Usp29* imprinted domain visualized at 25, 10, 5, and 2.5 kb resolutions. Paternal contacts shown in blue, Maternal contacts shown in red. Square root vanilla coverage (VC_SQRT) method was used for normalization. Matrices are visualized in the GRCm39/mm39 mouse annotations.

**Extended Figure 2.**
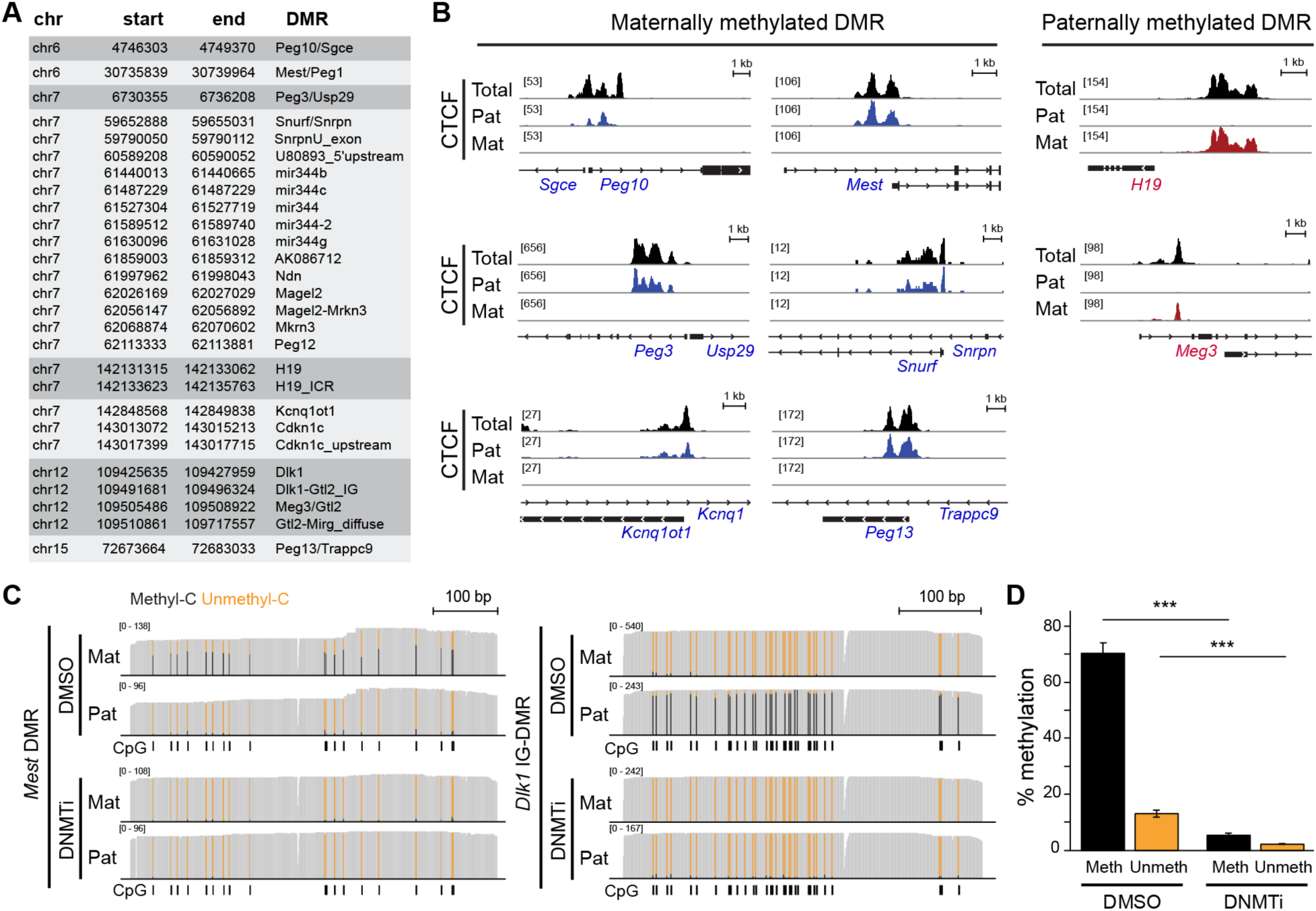
Methylation-sensitive CTCF binding at ICRs organizes allele-specific chromatin architecture. (**A**) All annotated DMRs in mouse cortex located within the capture regions. (**B**) Counts per million (CPM) normalized total and allele-specific CTCF tracks from primary cortical neurons at the indicated DMRs. (**C**) Representative nanopore sequencing tracks of PCR products derived from bisulfite-treated genomic DNA from DMSO and DNMTi-treated mESCs. Methylated CpG (black); unmethylated CpG (orange). (**D**) Average methylation percentage at CpGs quantified from (**C**). n=3 replicates per group. t-test *** < 0.001.

**Extended Figure 3.**
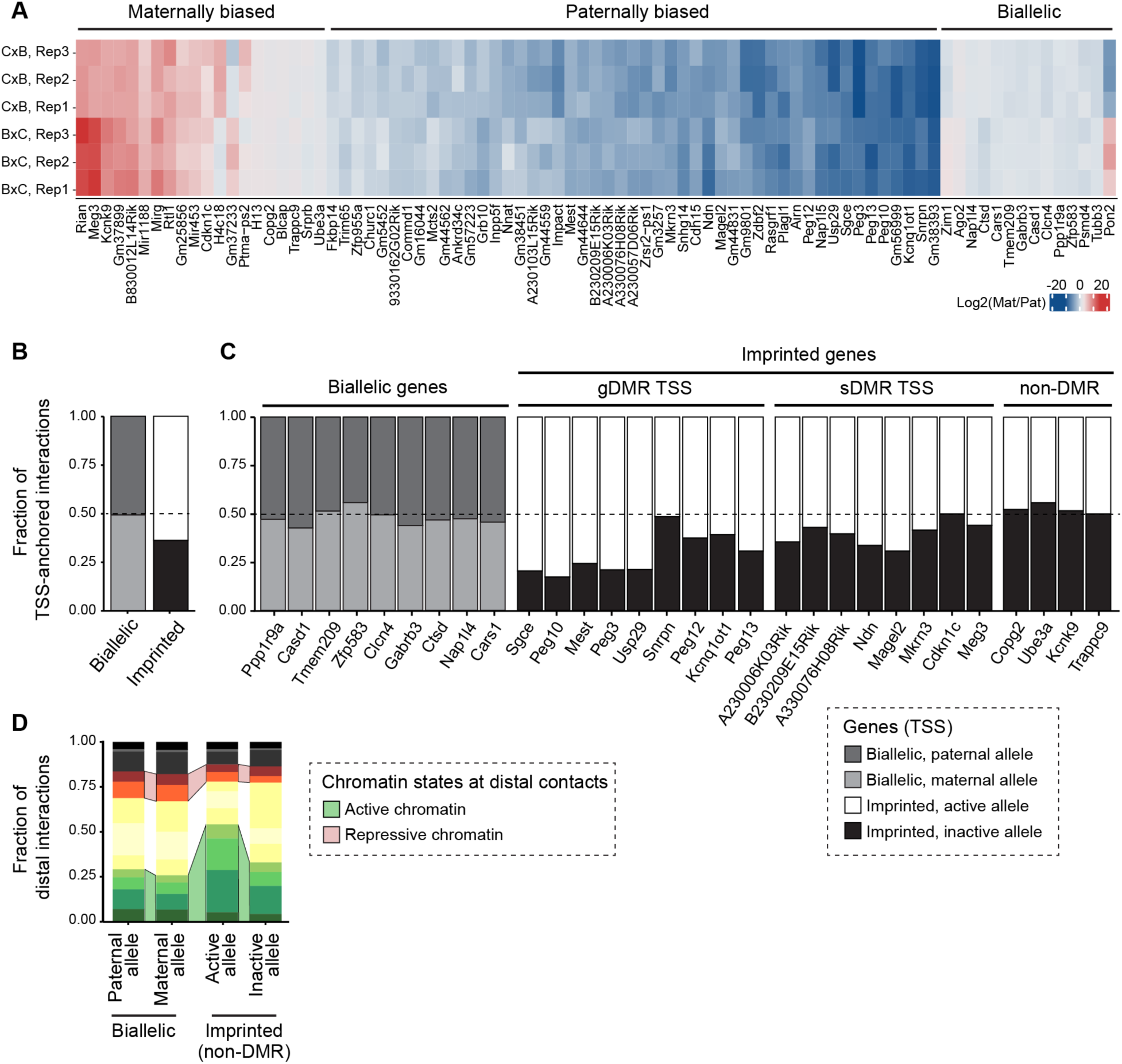
Imprinted gene promoters exhibit allelic differences in distal chromatin contacts in primary cortical neurons. (**A**) Heatmap of allele-specific RNA expression from primary cortical neurons. Rows correspond to replicates from BxC or CxB F1 neurons. Colors represent allelic expression biases. (**B**) Fraction of total TSS-anchored contacts from each allele of all biallelic or imprinted genes in the capture region from primary cortical neurons. (**C**) Fraction of TSS-anchored contacts from each allele of individual biallelic or imprinted gene in primary cortical neurons. Only a representative subset of biallelic genes is shown for illustrative purposes. (**D**) Fraction of TSS-anchored contacts colored by chromatin state of distal contact point. n = 4 replicates of primary cortical neuron capture Hi-C.

**Extended Figure 4.**
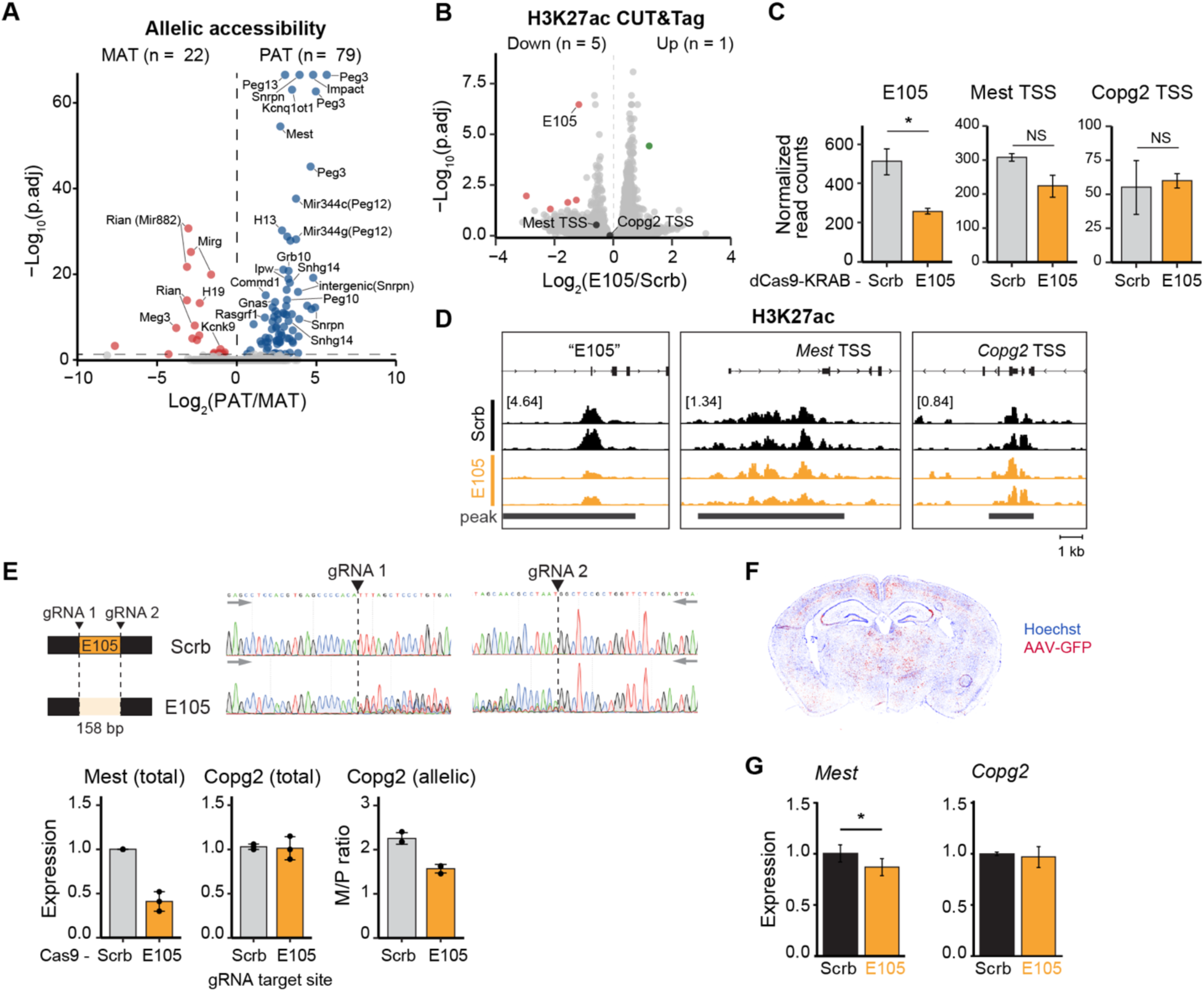
CRISPRi screening uncovers functional enhancers at imprinted domains. (**A**) Volcano plot of maternal vs paternal biased ATAC-seq peaks from mouse cortex. n = 3 replicates per genotype, 6 in total. (**B**) Volcano plot of differentially regulated H3K27ac peaks in differentiated NPCs upon E105 targeting gRNAs transduction. (**C**) Bar graphs showing the normalized read counts in Scrb vs E105 gRNA transduced samples. (**D**) H3K27ac CUT&Tag tracks upon transduction of Scrb vs E105 targeting gRNAs into dCas9-KRAB NPCs followed by differentiation. CUT&Tag peak called by SICER2 pipeline shown in black bar. (**E**) Validation of E105 function by Cas9-mediated enhancer deletion in NPCs followed by differentiation. *Top:* Sanger sequencing traces confirming enhancer deletion. *Bottom: Mest* and *Copg2* total (RT-qPCR) or allelic (ddPCR) expression. (**F**) Patched images of mouse brain section showing Hoechst 33342 straining, GFP signal derived from AAV, or merged. (**G**) Relative expression of *Mest* and *Copg2* in Scrb vs E105 targeting gRNA AAV-injected mouse cortex measured by qRT-PCR (n = 5 mice). t-test * < 0.05.

**Extended Figure 5.**
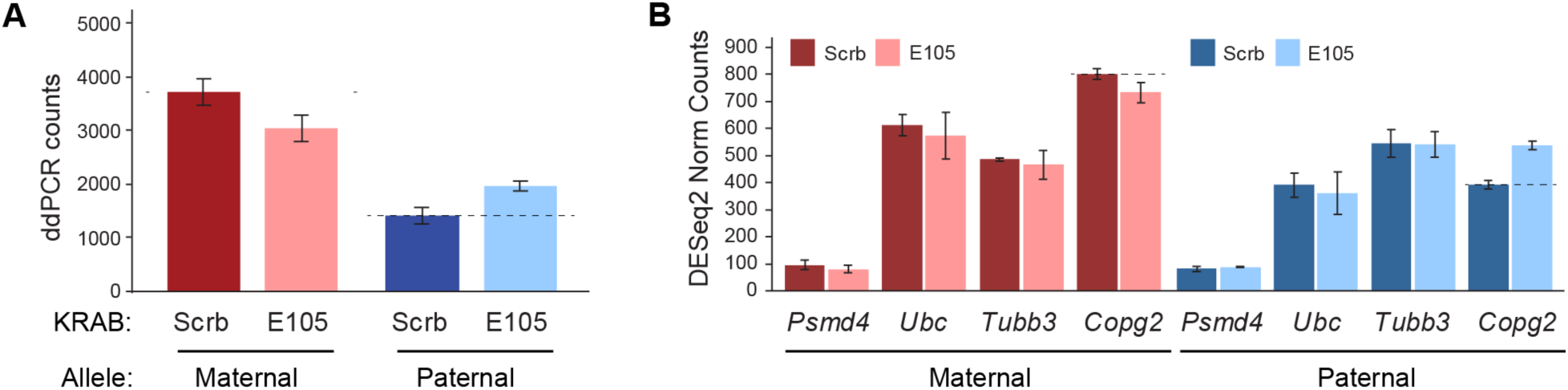
*Copg2* is regulated by both a distal enhancer and *Mest*-XL lncRNA. (**A**) Raw maternal vs paternal ddPCR counts showing cDNA copy numbers in Scrb vs E105 gRNA samples. (**B**) Maternal vs paternal normalized read counts in Scrb vs E105 gRNA transduced NPCs followed by differentiation.

